# The Promise of Deep Learning for BCIs: Classification of Motor Imagery EEG using Convolutional Neural Network

**DOI:** 10.1101/2021.06.18.448960

**Authors:** Navneet Tibrewal, Nikki Leeuwis, Maryam Alimardani

## Abstract

Motor Imagery (MI) is a mental process by which an individual rehearses body movements without actually performing physical actions. Motor Imagery Brain-Computer Interfaces (MI-BCIs) are AI-driven systems that capture brain activity patterns associated with this mental process and convert them into commands for external devices. Traditionally, MI-BCIs operate on Machine Learning (ML) algorithms, which require extensive signal processing and feature engineering to extract changes in sensorimotor rhythms (SMR). However, in recent years, Deep Learning (DL) models have gained popularity for EEG classification as they provide a solution for automatic extraction of spatio-temporal features in the signals. In this study, EEG signals from 54 subjects who performed a MI task of left-or right-hand grasp was employed to compare the performance of two MI-BCI classifiers; a ML approach vs. a DL approach. In the ML approach, Common Spatial Patterns (CSP) was used for feature extraction and then Linear Discriminant Analysis (LDA) model was employed for binary classification of the MI task. In the DL approach, a Convolutional Neural Network (CNN) model was constructed on the raw EEG signals. The mean classification accuracies achieved by the CNN and CSP+LDA models were 69.42% and 52.56%, respectively. Further analysis showed that the DL approach improved the classification accuracy for all subjects within the range of 2.37 to 28.28% and that the improvement was significantly stronger for low performers. Our findings show promise for employment of DL models in future MI-BCI systems, particularly for BCI inefficient users who are unable to produce desired sensorimotor patterns for conventional ML approaches.

## 1 Introduction

Motor Imagery (MI) is a dynamic experience where the user contemplates mental imagination of motor movement without activation of any muscle or peripheral nerve. A Motor Imagery Brain-Computer Interface (MI-BCI) serves as a system that converts brain signals generated during such imagination into an actionable sequence (Alimardani et al., 2018; Cho et al., 2018; Millán et al., 2010; Pfurtscheller & Neuper, 2001)

MI-BCI systems mainly utilize electroencephalogram (EEG) for measurement of brain activity (Lebedev & Nicolelis, 2017). EEG provides high temporal resolution, can be portable, is relatively low cost and represents synchronous electrical signals produced by the brain (Lebedev & Nicolelis, 2017). However, the recorded EEG signals are non-stationary and suffer from a low signal-to-noise ratio (SNR) and poor spatial resolution. Therefore, in order to employ them in a BCI system, it is necessary to apply advanced signal processing techniques to clean the data from artefacts and extract relevant spatial, temporal and frequency information from the signals for the classification problem (Bharne & Kapgate, 2014).

Traditionally, MI-BCIs operate on machine learning (ML) algorithms in which spatial features associated with movement imagination are recognized. The imagining of a left or right body movement is accompanied by a lateralization of event-related (de)synchronization (ERD/ERS) in the mu (7-13 Hz) and beta (13-30 Hz) frequency bands of EEG signals (Pfurtscheller et al., 2006; Avanzini et al., 2012; Barros & Neto, 2018; Wang et al., 2019). This brain activity feature serves as an input to the ML algorithm classifying the imagined body movements. Therefore, the system relies on the user to consciously modulate their brain activity such that the lateralization can be detected. It is shown that fifteen to thirty percent of users cannot accomplish distinctive brain waves such that the classifier reaches accuracy above 70%. This is called ‘*BCI illiteracy*’ (Allison & Neuper, 2010) or ‘*BCI inefficiency*’ (Thompson, 2019), where a user is considered unable to control a BCI, even after extensive training. But the issue of BCI inefficiency might be argued more nuanced, as successful BCI control depends on a synergy between man and machine, and therefore enhancements on both sides are needed to reach efficient control (Thompson, 2019).

In almost half of MI-BCI studies (Wierzgala et al., 2018), the mu suppression lateralization is picked up by the Common Spatial Pattern (CSP) algorithm that linearly transforms EEG data into a subspace with a lower dimension in which the variance of one class (the imagined side) is maximized while the variance of the other class is minimized (Khan et al., 2019; Shen et al., 2017). The output of the CSP filter is then used as an input for a ML algorithm, such as linear discriminant analysis (LDA), support vector machine (SVM), or logistic regression (LR) to distinguish EEG patterns associated with motor imageries (Miao et al., 2020). LDA is a very popular model for binary classification of the MI task (Yuksel & Olmez, 2015); it works on the concept of minimizing the ratio of within-class scatter to between-class scatter while keeping the intrinsic details of the data intact (Shashibala & Gawali, 2016). Hence, LDA creates a hyperplane in the feature space based on evaluation of the training data to maximize the distance between the two classes and minimize the variance of the same class (Aydemir & Kayikcioglu, 2013; Hasan et al., 2015).

Although, ML techniques are commonly used for binary classification of MI-BCIs systems, they are extremely vulnerable to variability between subjects and drifts in the brain signals (Millán et al, 2010). ML techniques do not work well under the influence of noise and outliers, which are difficult to segregate from the primary data (Müller et al., 2004). Additionally, the performance of ML classifiers is highly dependent on the type of feature extraction technique that is used (Hsu, 2010). More importantly, they suffer from the ‘*curse of dimensionality*’ and are therefore highly susceptible to overfitting (AlZoubi et al., 2008). The curse of dimensionality stems from an imbalance between the number of extracted features and the number of training EEG patterns (i.e. number of subjects). In order to extract relevant information from the EEG data, multiple feature extraction techniques are adopted, which add more and more dimensions to the feature space (AlZoubi et al., 2008; Lotte et al., 2018). This creates a situation in which the features vastly outnumber the observations, resulting in overfitting and an erroneous model performance. Therefore, ML approaches require yet another step of feature selection for reduction of dimensionality in the training data, which yields additional computational costs in terms of memory usage and CPU time.

Deep Learning (DL) classifiers are a promising alternative to address the complexity of EEG signals, as they can work with raw data and directly learn features and capture structure of a large dataset without any feature engineering or selection processes (Albawi et al., 2018; Robinson et al., 2019; Wang et al., 2018; Yang et al., 2015). Thus, the issue of information loss while generating and selecting features is avoided when DL classifiers are used (Qiao & Bi, 2019). Additionally, they can be used to stabilize the learning process by overcoming the issue of noise and outliers in the data (Al-Ani et al., 2010). DL generates high-level abstract features from low-level features by identifying distributed patterns in the acquired data. Hence, DL models hold the potential of handling complex and non-linear high dimensional data (Wang et al., 2019).

Past research has already established the effectiveness of the DL approach, especially Convolutional Neural Network (CNN), in classification of MI-EEG (Tang et al., 2017; Gao et al., 2018; Sakhavi et al., 2015; Li et al., 2020; Dai et al., 2019; Tayeb et al., 2019; Stieger et al., 2020; Zhang et al., 2021; Ko et al., 2020; Mane et al., 2020). The advantages of CNN model include handling raw data without any feature engineering process, facilitating end-to-end learning and requiring lesser parameters than other deep neural networks (Shen et al., 2017; Albawi et al., 2018). CNN works well with large datasets and can exploit the hierarchical structure in natural signals (Schirrmeister et al., 2017). Moreover, CNN has good regularization and degree of translation invariance properties along with the ability to capture spatial and temporal dependencies of EEG signals (Aggarwal & Chugh, 2019). CNN can be particularly useful in classification of MI-EEG for low aptitude users. Stieger et al. (2020) showed a negative correlation between online (ML-based) performance and improvement of accuracy with CNN, which suggests that BCI inefficient users may benefit from applying a DL classifier, even more than high aptitude users. They further showed that the low performing users in the online classification did not necessarily produce the expected SMR activity during MI process, but instead produced differentiating activity over brain regions outside the motor cortex such as occipital and frontal gamma power, which could not be recognized by CSP. Therefore, DL methods might be beneficial in improving performance for inefficient users and serve as a promising tool in enhancing overall BCI usability.

This study aims to compare the two approaches of ML and DL in classification of MI EEG signals in a large group of 54 subjects. In most of previous studies, CNN has been compared with ML classifiers other than CSP+LDA. However, the use of CSP+LDA model is widespread in binary MI-BCI classification (Lotte et al., 2018; Nicolas-Alonso & Gomez-Gil, 2012; Selim et al., 2018). Hence, in this study, for every subject, a CNN model (DL approach) was trained and its performance was compared with the conventional CSP+LDA model (ML approach).

Figure 1 shows sequential steps that were taken in each approach to construct a MI-BCI classifier and obtain classification performances. The ‘*Signal Acquisition*’ step was carried out through EEG to monitor the brain signals arising from the mental image of the movement by the user. The complexity of the ML approach arises with the steps involved in ‘*Pre-processing*’ and ‘*Feature Extraction*,’ whereas in the DL approach, raw data can directly be fed into the model. Hence, by applying both approaches to the data from 54 subjects, this study intends to answer the following research question: “*Can a CNN classifier trained with raw EEG signals achieve a higher performance than a machine learning model that runs on processed EEG features for classification of a two-class Motor Imagery task?*”

**FIGURE 1.**
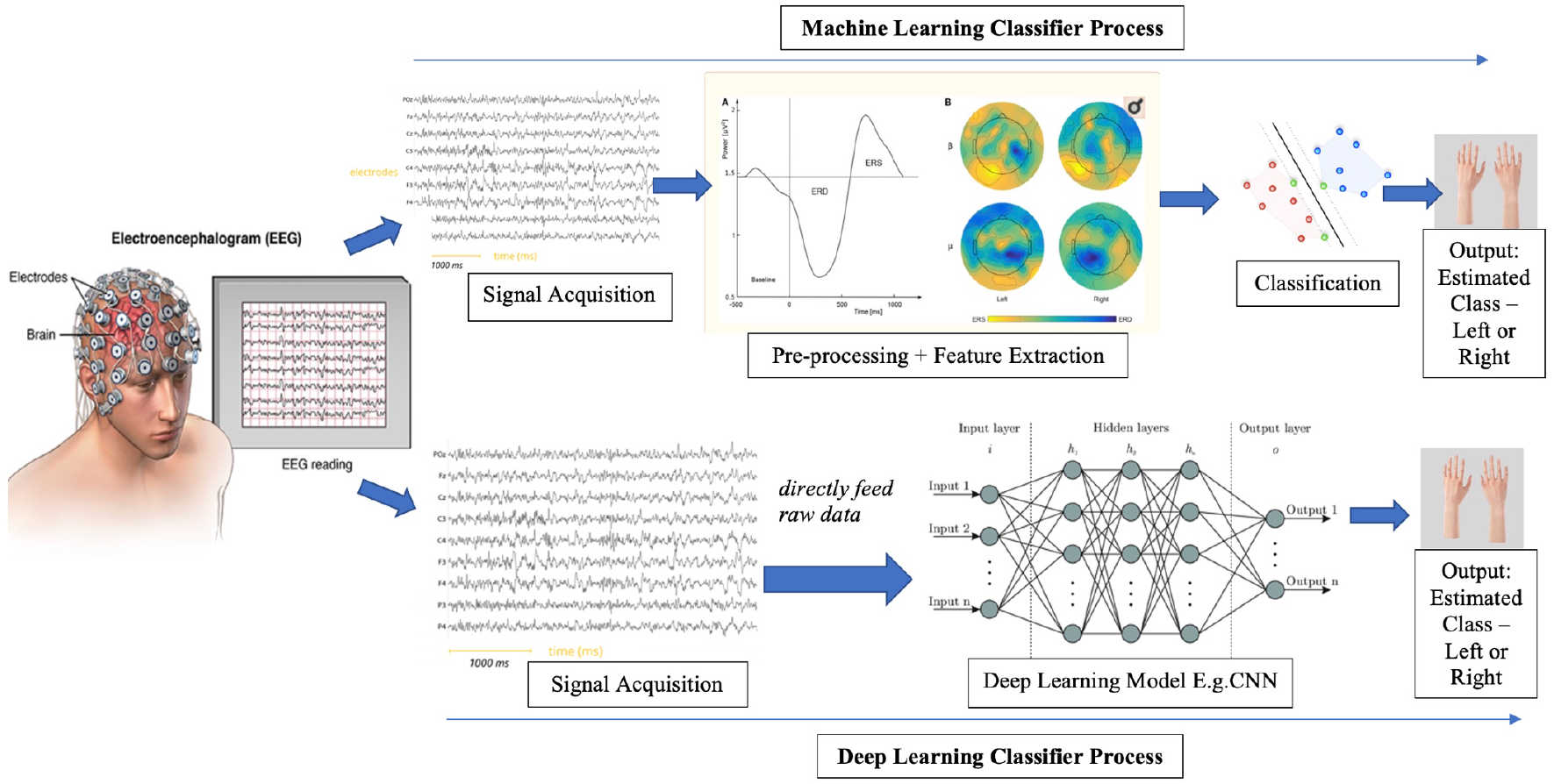
An overview of MI-BCI classification using machine learning vs. deep learning approaches. In ML approach, EEG signals are first pre-processed and relevant features are extracted before applying a classifier. In DL approach, raw signals are directly fed into the model.

## 2 Methods

In order to compare conventional ML models with a DL approach in a large group of novice BCI users, EEG signals were collected from 57 subjects while they performed the MI task using an existing BCI system. Thereon, the recorded EEG signals were used to train a CNN and CSP+LDA model to conduct an offline classification of two-class MI task. The following section gives a description of the data collection procedure and details of the classification models.

### 2.1 Experiment

#### 2.1.1 Participants

In this experiment, 57 subjects participated (21 male, 36 female, *M*_*age*_ = 20.71, *SD*_*age*_ = 3.52). All of them were right-handed and novice to BCI and the MI task. The Research Ethics Committee of Tilburg School of Humanities and Digital Sciences approved the study (REDC #20201003). All subjects received explanation regarding experiment procedure and signed a consent form before the experiment.

#### 2.1.2 EEG Acquisition

Sixteen electrodes recorded EEG signals from the sensorimotor area according to the 10-20 international system (F3, Fz, F4, FC1, FC5, FC2, FC6, C3, Cz, C4, CP1, CP5, CP2, CP6, T7, T8). The right earlobe was used as a reference electrode and a ground electrode was set on AFz. Conductive gel was applied to keep the impedance of the electrodes below 50 kOhm. Subjects were instructed to sit calmly and avoid movements and excessive blinking. The signals were amplified by a g.Nautilus amplifier (g.tec Medical Engineering, Austria). The data was sampled at 250 samples/second. The noise during EEG recording was reduced by applying a 48-52 Hz notch filter and 0.5-30 Hz bandpass filter.

#### 2.1.3 Motor Imagery Task

Participants performed the MI task in four runs, each consisting of 20 right-and twenty left-hand trials. The first run was a non-feedback run, followed by three runs in which the subjects received feedback in form of a feedback bar on the computer screen. The feedback bar presented the classification certainty as computed by the g.tec BCI classifier, which relies on the CSP+LDA approach. The classifier was calibrated for each subject based on the data of the latest run while the subject took a break between the runs.

In total, participants performed 120 MI trials. Each MI trial took eight seconds. The timeline of each trial is shown in Figure 2. It started with a fixation cross that was displayed in the center of the screen for three seconds. In the next 1.25 seconds, a red arrow cued the direction of the trial; the subject had to imagine squeezing their left hand if the arrow pointed to the left and their right hand if the arrow pointed to the right, without tensing their muscles. During the last 3.75 seconds, the calibration run showed the fixation cross again (see Figure 2a), while the feedback runs showed a blue feedback-bar indicating the direction and certainty of the algorithms’ classification (see Figure 2b). Participants were instructed to stay focused on the imagination of the movement even during the feedback and try to not get distracted by it. The end of the trial was marked by a blank screen. The rest time between trials varied randomly between 0.5 and 2.5 seconds.

**FIGURE 2.**
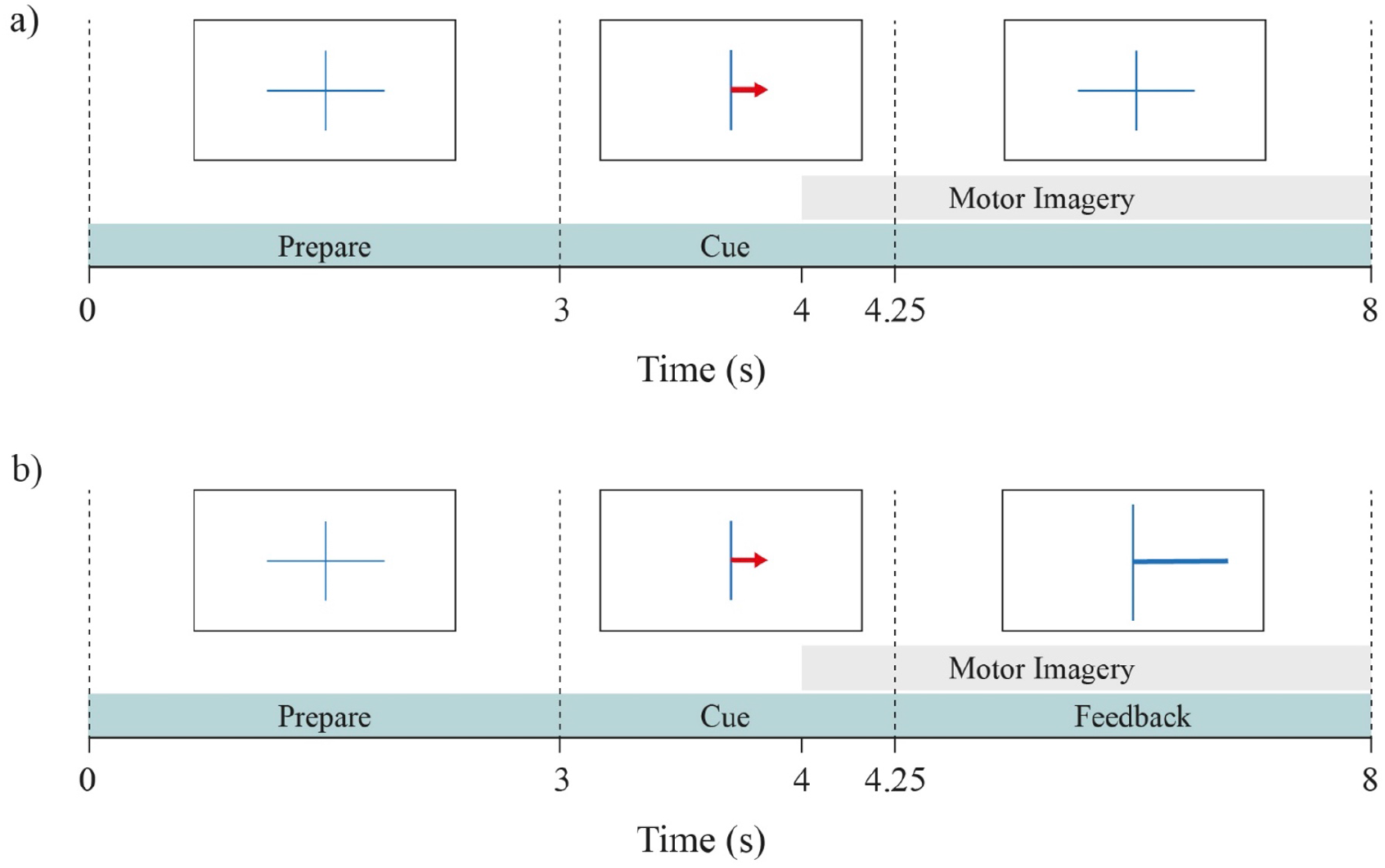
The time course of each trial in the BCI task. (a) shows the calibration run and (b) the feedback runs. In all trials, participants saw a fixation cross and thereafter an arrow pointing to either left or right, which indicated the corresponding hand for the MI task in the trial. In feedback runs, the blue bar indicated the direction and certainty of the classifier’s prediction in order to feedback to the participants. The grey area indicates the time course of the MI task.

#### 2.1.4 EEG Dataset

The signals from three participants were not recorded in a satisfactory manner due to technical issues during the experiment. Hence, only 54 participants were chosen from the dataset for this study. An epoch of 4 seconds was selected from each trial. This epoch, targeting the MI period, started at second 4 of the trial (1 second after cue presentation) and ended at second 8 (5 seconds after cue presentation), which is in line with the study of Marchesotti et al. (2016). The selected time segment is indicated with the grey area in Figure 2.

### 2.2 Machine Learning Model

The ML approach consisted of preprocessing the signals, constructing CSP filters for feature extraction and an LDA model for classification of the left vs. right classes. CSP is a feature extraction technique that selects spatial filters from multi-channel signals and then linearly transforms EEG data into a subspace with lower dimension that maximizes the variance of one class while minimizing the variance of the other class (Khan et al., 2019; Shen et al., 2017). CSP algorithm is widely used in binary MI-BCIs due to its computational simplicity and improving signal to noise ratio (Bashashati et al., 2015; Guan et al., 2019). The output of CSP can be used as input for the LDA classifier in order to distinguish the classes of MI task.

LDA is a dimensionality reduction model that works on the concept of minimizing the ratio of within-class scatter to between-class scatter while keeping the intrinsic details of the data intact (Shashibala & Gawali, 2016). Hence, LDA creates a hyperplane in the feature space based on evaluation of the training data to maximize the distance between the two classes and minimize the variance of the same class (Aydemir & Kayikcioglu, 2013; Hasan et al., 2015). LDA is very popular for binary classification of the MI task (Yuksel & Olmez, 2015).

#### 2.2.1 Architecture

Before applying the ML model, the EEG signals recorded from the participants were pre-processed and temporally filtered to remove artifacts. Data containing bad impedance, error in recording, or excessive movement-related noise were removed (3 subjects, see 2.1.4). Then the EEG signals corresponding to the onset of MI task (second 4 to 8, see Figure 2) were selected and taken into account (Park & Chung, 2019). Thereon, Filter Bank Common Spatial Pattern (FBCSP) was used to extract subject-specific frequency band of 7-30 Hz from the data through the implementation of fifth order Butterworth (Park & Chung, 2019; Lotte & Guan, 2011).

FBCSP was used because it is instrumental in discriminating the binary classification of EEG measurements (Ang et al., 2012; Raza et al., 2015; Park & Chung, 2019). It should be noted that CSP is highly dependent on the selection of frequency bands, however there is no optimal solution to select the right filter bank (Kumar et al., 2017). Using a filter bank before CSP helps to improve the accuracy level of the model (Yahya et al., 2019). A wide range of 7-30 Hz is usually adopted for CSP when used for MI classification (Kumar et al., 2017). Hence, the frequency bandwidth was kept between 7-30 Hz covering the mu and beta bands that are required to analyze Event-Related Desynchronization (ERD) and Event-Related Synchronization (ERS) from the MI brain signals.

After the pre-processing and filtering steps, the 120 MI trials of each participant were concatenated and randomized. CSP algorithm was performed on each participant’s data using the ‘*scikit*’ package in Python (Yuksel & Olmez, 2015). CSP extracted the spatially distributed information from the output of FBCSP by linearly transforming the EEG measurements in order to define discriminative ERD/ERS features (Ang et al., 2012; Park & Chung, 2019; Raza et al., 2015). Once feature extraction was completed, ‘*scikit*’ package was again used to implement the LDA classifier in order to reduce the dimensionality of the sub-bands and to perform binary classification (Vidaurre et al., 2011).

### 2.3 Deep Learning Model

The DL model was constructed by feeding raw EEG signals directly into a CNN model.

CNN is a feed-forward Artificial Neural Network (ANN) model and has a sequence of layers where every layer is the output of an activation using a differential function (Aggarwal & Chugh, 2019). In a CNN, the inputs are assembled to different layers of neurons, each representing a linear combination of the inputs (Pérez-Zapata, 2019). The learning process involves modification of the parameters by adjusting weights between different layers in order to achieve the desired output (Roy et al., 2019). The learning continues until the training set reaches a steady state where the weights become consistent and an optimal output is reached (Roy et al., 2019). During the training phase of the CNN model, different layers can extract features at a different level of abstraction (Roy et al., 2019). The initial layers learn local features from the raw input, and the end layers learn global features (Schirrmeister et al., 2017).

#### 2.3.1 Architecture

A 2D CNN model was constructed using ‘*keras*’, a high-level neural networks API written in Python (Keras, 2019). Figure 3 shows the architecture of the proposed CNN model.

**FIGURE 3.**
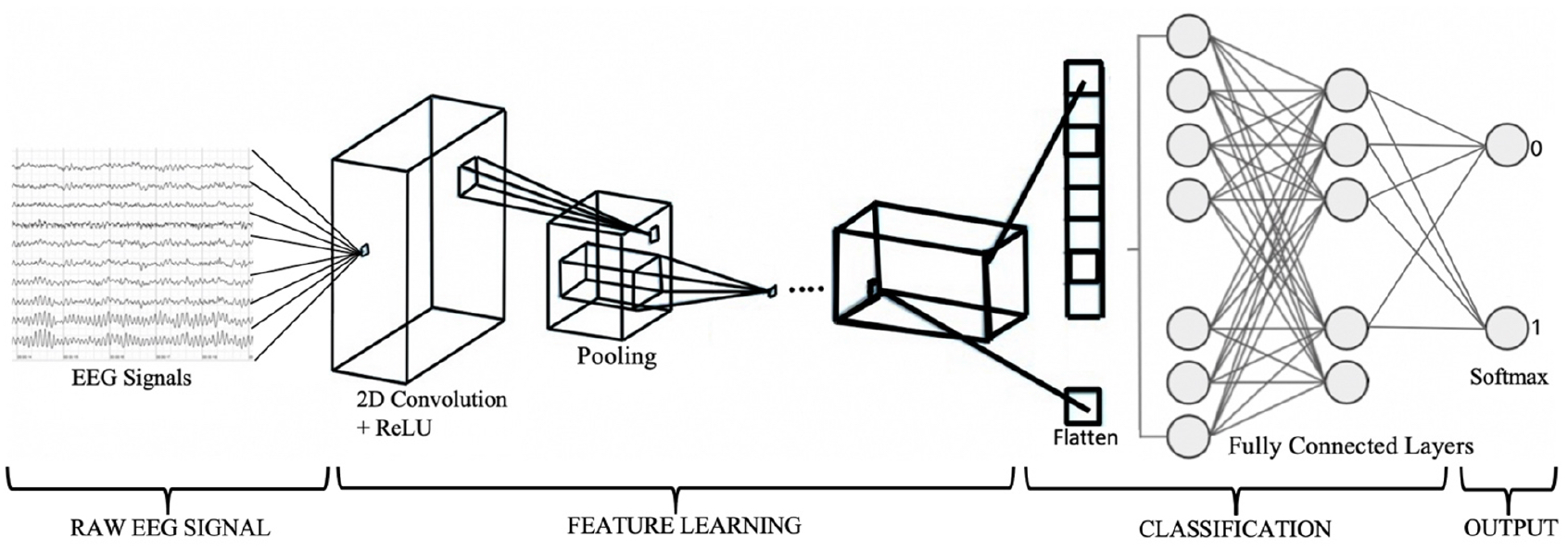
CNN Architecture.

The first two components of the architecture are the number of convolution filters used and the kernel size that specifies the height (columns) and width (rows) of the 2D convolution window. These were set to 30 and 5×5 respectively. The dimensions of the input shape applied were (1×4×4). In order to compute a network’s hidden layers, activation functions should be implemented (Goodfellow et al., 2016). For this task, Rectified Linear Function (ReLU) was used. ReLU conducts simple mathematical operations, preserves characteristics that result in good generalization and is less computationally expensive than other approaches. Moreover, ReLU has the advantage of the speed and overcoming gradient leakage issue when compared with other activation functions (Pérez-Zapata, 2019).

Max pooling was added to the model in order to downsample the input and refrain from losing important data features. The size of 2×2 was used based on the works of Dharamsi et al. (2017) and Abbas and Khan (2018). The output of max pooling was flattened into a vector of input data by executing a flatten layer (Goodfellow et al., 2016). Subsequently, three dense layers were added. The first two implemented a linear function in which all inputs were connected to all outputs by a specific weight (Ullah et al., 2019). The units of these were set to 256 and 128 and were activated by ReLU functions. The final dense layer’s units were fixed to 2 as this was the number of class labels in the data. Finally, Softmax was applied to the last (output) layer as an activation function, used for class classification tasks (Goodfellow et al., 2016).

#### 2.3.2 CNN Model Compilation

The hyperparameters implemented in the 2D CNN model’s compilation phase are the loss function, the optimizer and the evaluation metric. Since the dataset has two target labels (right and left), the loss function categorical cross-entropy was applied. The optimizer *‘Adam’* was used because it is a widely used gradient-based optimization of stochastic objective functions (Kingma & Ba, 2014). An essential parameter of *‘Adam’* is the learning rate, which regulates the modification of the model based on the error obtained from the updated weights (Kingma & Ba, 2014). For the task at hand, the learning rate was set to its default value of 0.01. The evaluation metric was set to accuracy to delineate how well the CNN model could classify left vs. right MI EEGs. (Goodfellow et al., 2016).

#### 2.3.3 CNN Model Fit

During model fitting, a specified batch size and number of epochs need to be adopted for backpropagation to take place (Browniee, 2016). The batch size greatly influences the time to converge and the amount of overfitting (Radiuk, 2018); a big batch takes into account many samples to calculate a gradient step and therefore might slow down the model training (Goodfellow et al., 2016). On the other hand, small batch sizes can supervise variation in the distribution. The batch size for the 2D CNN model was set to 264.

An epoch in DL means that all the samples in the training set are traversing through the model once (Browniee, 2016). This helps the network to see previous data for readjusting the model parameters in order to reduce any biases. The neural network updates the weights of the neuron during each epoch (Torres, 2018). However, there is not any prescribed method to calculate how many epochs are required for a particular model. Sharma (2017) stated that different values of epochs should be tried until the learning curve of the model moves from underfitting to an optimum level and until overfitting attributes start showing up, then the subsequent epoch size should be deemed as the threshold for the model. Thus, as long as both training and test accuracies are increasing at an equivalent rate, the training of the model should continue (TensorFlow, 2020). Considering the arguments from Kingma and Ba (2015) and TensorFlow (2020), 500 epochs per subject was deemed to be the threshold for the CNN model.

### 2.4 Evaluation

For the CNN model, the data was split into 80% training and 20% test data. Tang et al. (2017) used the same splitting variation for building their CNN model. Accuracy is defined as the total amount of correct predictions that the model made including both training and test accuracies (Goodfellow et al., 2016). Hereby, the mean accuracy over all the subjects in training and test phase was calculated in order to compare the performance of CSP+LDA and CNN models.

Additionally, we observed how the CNN model and CSP+LDA model performed subject-wise by computing the difference of the two models’ accuracy for each subject. This was done to give greater validity to the findings as inter-subject variability can affect the overall performance of a classifier (Saha & Baumert, 2020). While accuracy is the overall evaluation measure of a model, it does not fully exhibit its prediction capacity. Therefore, in addition to the overall prediction accuracy, we extracted F-score metric for each class of *‘left’* or *‘right’* MI. F-score is the harmonic mean of the precision and recall metrics and demonstrates the discriminant power of the model for each existing class in the data. Previous research has shown that the BCI user handedness plays a role in lateralization of ERD/ERS during the MI task (Zapala et al., 2020). In our study, all subjects were right-handed, therefore it was expected that the errors made by the model would be more for one MI class than the other.

## 3 Results

The average score of the training and test accuracies across 54 subjects were taken into consideration to report the performance level of the CNN and CSP+LDA models. The CNN model reached an average training accuracy of 80.58% (*SD* = 5.01) and an average test accuracy of 69.42% (*SD* = 4.97), whereas the average training and test accuracies for the CSP+LDA model were 52.54% (*SD* = 5.12) and 52.56% (*SD* = 2.08), respectively. Table 1 gives a summary of these results.

**TABLE 1.** Comparison between training and test accuracies of CNN and CSP+LDA models.

| Model | Training Accuracy<br>($N=54$) | | Test Accuracy<br>($N=54$) | |
| --- | --- | --- | --- | --- |
| | Mean | $SD$ | Mean | $SD$ |
| CNN | 80.58 | 5.01 | 69.42 | 4.97 |
| CSP+LDA | 52.54 | 5.12 | 52.56 | 2.08 |

The obtained accuracies for both CNN and CSP+LDA models were normally distributed as evaluated with Shapiro-Wilk test (CNN: *W* = 0.98, *p* = .66; CSP+LDA: *W* = 0.97, *p* = .12). Therefore, a pairwise t-test was employed to compare the test accuracies obtained from the DL classification method to those of the ML approach (*t*(53) = 22.12, *p* < .001). This indicated that the CNN classifier significantly outperformed the CSP+LDA approach by 15.32 to 18.38% within the 95% confidence interval.

Table 2 contains the top ten accuracy rates observed in the subjects using the CNN and the CSP+LDA model. As it can be seen in this table, the highest accuracy achieved by the CNN model for a subject was 81.80%, whereas the CSP+LDA model could only reach a highest accuracy rate of 57.17%. Also, although not included in this table, it was observed that the lowest accuracy level obtained from the CNN model across all the subjects was 58.60%, which is still higher than the highest accuracy rate obtained by the CSP+LDA model across all the subjects.

**TABLE 2.** Top ten highest classification accuracies achieved by the CNN model and the CSP+LDA model.

| CNN Model |  | CSP + LDA Model |  |
| --- | --- | --- | --- |
| Participant ID | Accuracy % | Participant ID | Accuracy % |
| Subject 31 | 81.80 | Subject 29 | 57.17 |
| Subject 48 | 77.36 | Subject 7 | 56.70 |
| Subject 16 | 76.94 | Subject 55 | 56.41 |
| Subject 25 | 76.32 | Subject 54 | 56.23 |
| Subject 55 | 75.81 | Subject 34 | 56.16 |
| Subject 9 | 75.72 | Subject 8 | 55.90 |
| Subject 60 | 75.64 | Subject 30 | 55.67 |
| Subject 12 | 75.58 | Subject 32 | 55.65 |
| Subject 42 | 75.52 | Subject 60 | 55.32 |
| Subject 23 | 75.08 | Subject 66 | 54.50 |

To obtain an estimation of the subject-wise performance difference between the two models, the difference of the obtained accuracy from the CNN model and the CSP+LDA model for each subject *(Accu*_*CNN*_ *– Accu*_*CSP+LDA*_) was computed. This subject-wise comparison revealed that the DL approach achieved a higher accuracy level for all subjects with a minimal difference of 2.37% and maximal difference of 28.28%. Figure 4 illustrates the number of subjects for whom the CNN model showed accuracy improvement in 6 bins of 1-5%, 6-10%, 11-15%, 16-20%, 21-25% and 26-30%. From this figure, it can be inferred that the CNN model outperformed the CSP+LDA model by more than 11% accuracy for 92.59% of the participants. Therefore, it can be concluded that CNN was able to extract intrinsic features from the EEG signals and thereon, performed classification with higher accuracy level.

**FIGURE 4.**
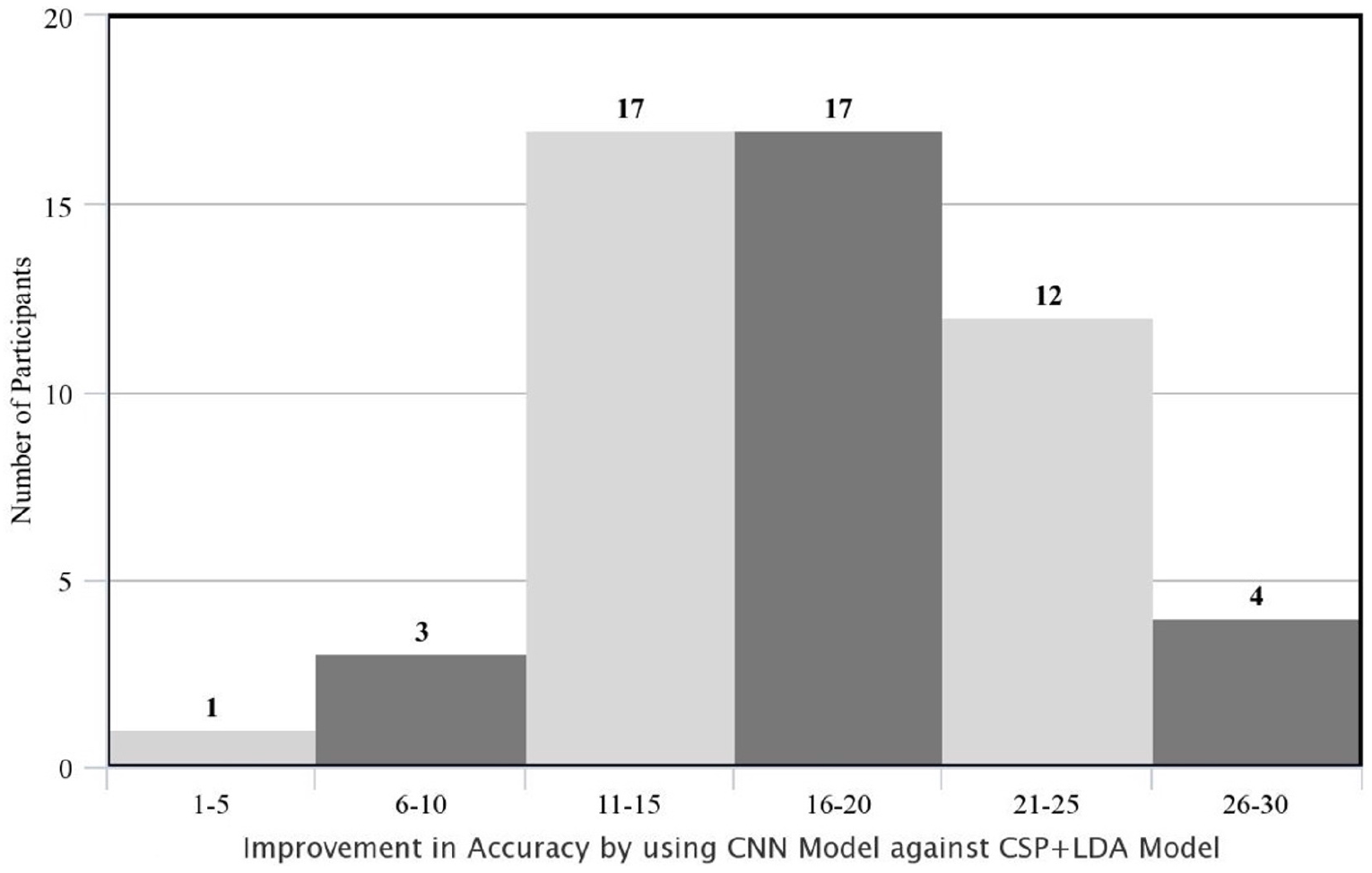
Improvement in the accuracy rate of the subjects using CNN model against CSP+LDA in percent points (i.e., absolute difference between the two accuracies; AccuCNN – AccuCSP+LDA).

Further exploration was done to investigate whether the improvement achieved by the CNN model was different across BCI users based on their initial MI skill. Traditionally, users that cannot produce desired ERD/ERS patterns to be recognized by a MI-BCI classifier are defined as low aptitude users or BCI inefficients (Thompson, 2019). Therefore, based on classification accuracy rates obtained from the CSP+LDA model, subjects were divided into two groups of Low Performers and High Performers. The split was made based on the accuracy median (*Med* = 52.14%), resulting in 27 subjects per group. For each group, the improvement of classification performance from the CSP+LDA model to the CNN model was obtained per subject by subtracting the model accuracies *(ΔAccu=Accu*_*CNN*_ *– Accu*_*CSP+LDA*_).

Figure 5 shows the mean accuracy improvement (*ΔAccu*) for each group. On average, the CNN model increased the accuracy rate of the Low Performers by 18.46% (*SD* = 4.98%) and the High Performers by 15.25% (*SD* = 5.81%). The obtained *ΔAccu* values for both Low Performer and High Performer groups were normally distributed as evaluated with Shapiro-Wilk test (Low Performers: *W* = 0.96, *p* = .47; High Performers: *W* = 0.98, *p* = .84). Therefore, an independent t-test was employed to compare them, revealing a significantly higher improvement of classification performance by the CNN model for Low Performers (*t*(26) = 2.18, *p* < .05). This result supports the notion that the CNN model can better capture intrinsic oscillation patterns associated with the MI task in inefficient BCI users, whose modulation of SMR cannot be successfully recognized by the CSP+LDA model.

**FIGURE 5.**
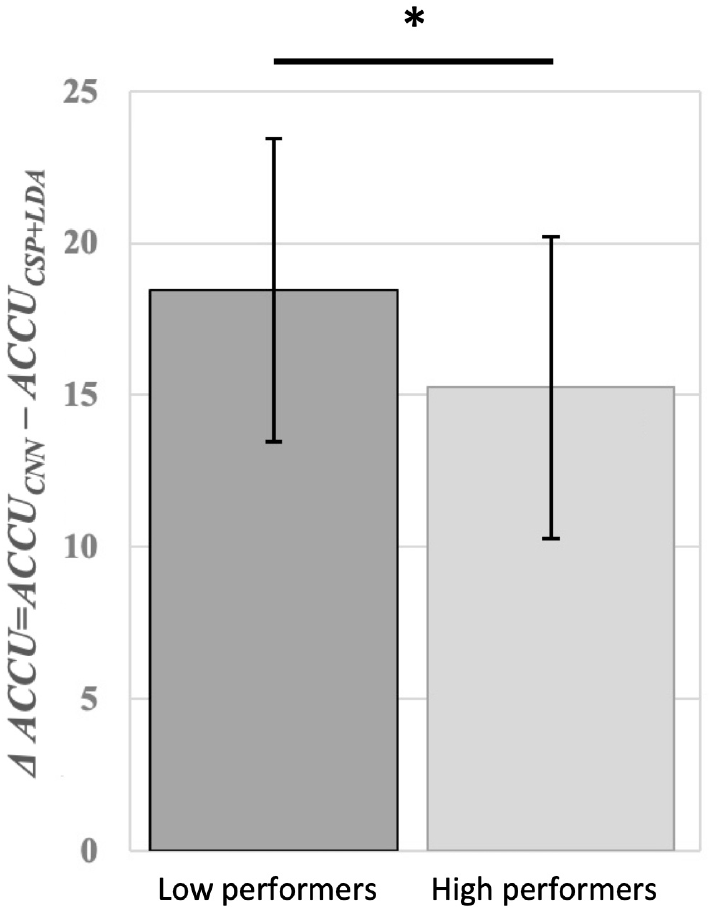
Mean difference between accuracies of CNN and CSP+LDA models (*Accu*_*CNN*_ *– Accu*_*CSP+LDA*_) for Low Performer and High Performer groups. Low Performers showed significantly higher improvement in MI-BCI accuracy after using a CNN classifier.

Finally, F-Score was calculated for each class in order to measure the predictive power of the classifiers with respect to the ‘*left*’ or ‘*right*’ MI movements. Table 3 summarizes the average and SD of F-Scores across all subjects obtained by the CNN and CSP+LDA models in regard to each MI class. As can be seen in this table, the CNN model achieved higher F-Score values for both ‘*left*’ and ‘*right*’ hand prediction compared to the CSP+LDA model. A pairwise t-test comparing the F-Scores of the two models found a significant difference for both ‘*left*’ MI movements (*t*(53) = 18.28, *p* < .05) as well as ‘*right*’ MI movements (*t*(53) = 19.47, *p* < .05) favoring CNN as a classifier beyond CSP+LDA approach.

**TABLE 3.**
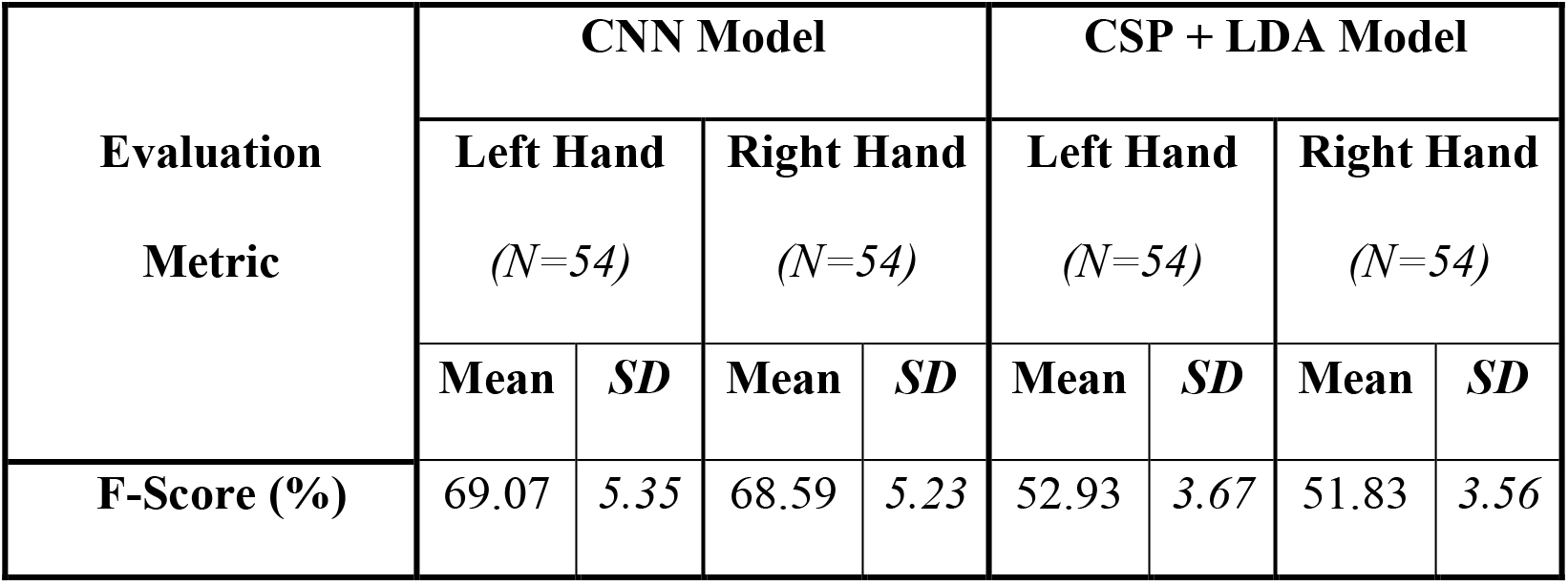
Average F-score obtained by the CNN and CSP+LDA models for each MI class.

| Evaluation<br>Metric | CNN Model |  |  |  | CSP + LDA Model |  |  |  |
| --- | --- | --- | --- | --- | --- | --- | --- | --- |
|  | Left Hand<br>(*N*=54) |  | Right Hand<br>(*N*=54) |  | Left Hand<br>(*N*=54) |  | Right Hand<br>(*N*=54) |  |
|  | Mean | *SD* | Mean | *SD* | Mean | *SD* | Mean | *SD* |
| **F-Score (%)** | 69.07 | 5.35 | 68.59 | 5.23 | 52.93 | 3.67 | 51.83 | 3.56 |

## 4 Discussion

In order for a BCI system to operate optimally for all users, it is crucial to devise a classification model that can learn from each individual’s brain signals and recognize task-related patterns with high accuracy. In this research, a CNN model was developed on a large EEG dataset from 54 subjects who conducted MI task during a BCI interaction. The main goal of this study was to validate that a DL approach employing raw EEG signals could outperform the state-of-the-art MI-BCIs, which often employ ML approach including CSP algorithm for feature extraction and LDA model for classification. Our results supported this hypothesis; the CNN model displayed significantly higher classification accuracy for the MI task as compared to the CSP+LDA approach for all users, but especially benefited low aptitude users by increasing their BCI performance significantly more than high aptitude users. Our results put forward the design of future BCI classifiers that facilitate better interaction between the user and the BCI system.

Until now, an in-depth analysis of a CNN model in which a large and novel dataset of raw EEG signals were directly fed to the model for classification of the MI task was still missing. Previous studies mainly focused on comparing different ML and DL models on already existing datasets. For instance, Sakhavi et al. (2015) employed the BCI competition IV (Dataset 2b) for multi-class classification of MI task. Their CNN model achieved accuracy level of 69.56%, whereas their Support Vector Machine (SVM) model, Multi Layer Perceptron (MLP) model and CNN+MLP model achieved accuracy level of 67.01%, 65.78% and 70.60%, respectively. Likewise, Li et al. (2020) used BCI competition IV (Dataset 2b) but the authors combined different feature extraction techniques with their ML and DL models to conduct comparison between these models. Li et al. (2020) showed that a combination of Continuous Wavelet Transform (CWT) with Simplified Convolutional Neural Network (SCNN) model achieved an average accuracy of 83%, which was 7.22%, 9.62%, 10.93%, 7.49%, 6.94%, 5.58% and 5.05 % higher than CNN+Stacked AutoEncoders (SAE), CSP, Adaptive Common Spatial Pattern (ACSP), Deep Belief Network (DBN), CSP+SCNN, Fourier Transform (FFT)+SCNN and Short Time Fourier transform (STFT)+SCNN, respectively. In another study by Gao et al. (2018), CSP was used for feature extraction and the CNN model was combined with Sparse Representation-based Classification (SRC) algorithm for binary classification of the MI task. The dataset adopted by Gao et al. (2018) was BCI competition III (Dataset IVa). Here the authors showed that their SRC+CNN model achieved mean accuracy of 80% (Gao et al., 2018).

Although previous studies provided promising results with a DL approach, the employed dataset by Sakhavi et al. (2015) and Li et al. (2020) only included nine subjects and the dataset used by Gao et al. (2018) only had five subjects. These datasets do not sufficiently represent the large inter-subject variability that exist among users (Leeuwis & Alimardani, 2020), which could affect the performance of the classifier. Different BCI users have a different state of mind, and hence different spatial, spectral and temporal patterns in their EEG signals (Ahn & Jun, 2015). Such variations can be due to the difference in the concentration levels of the participants while performing the MI task or baseline cognitive and psychological abilities (Leeuwis et al., 2021). Thus, it is necessary to perform BCI studies over a diverse and large pool of subjects in order to establish the broad generalizability of the findings. In comparison to previous studies that only employed datasets with limited number of participants and trials, this study collected MI EEG signals from 54 participants in three runs (120 trials) and built a 2D CNN model on our dataset. The large number of subjects in this dataset enabled us to statistically compare the subject-wise performance achieved by the CNN model as compared to the conventional ML approach. The results showed that the CNN model achieved an average of 69.42% accuracy across all subjects, which is similar to the CNN accuracy rate achieved by Sakhavi et al. (2015) who used feature engineering techniques to enhance the performance of their CNN model. The accuracy level achieved by this study might initially seem insufficient when compared to Gao et al. (2018) and Li et al. (2020), however, this difference can be explained by various pre-processing and feature engineering techniques that were employed by these two studies. Unlike past research, this study focused on evaluating the performance of CNN model without implementing any fine-tuning techniques and by directly feeding raw data into the model. The motive for this approach was to show the efficacy of deep learning models in exploiting information from raw data without any need for feature extraction. This makes deep learning models computationally more effective by eliminating the costly steps of pre-processing and feature extraction. Additionally, such neural networks can handle noise in EEG signals better than ML models and thus can provide a more robust performance in real-time BCI applications.

The low performance obtained in the ML approach has to be compared to the online classification accuracies presented in Leeuwis et al. (2021), where the average classification accuracy was 74.17%. This could be explained by different architectures: The online classification of Leeuwis et al. (2021) was conducted by g.BSanalyze software (g.tec Medical Engineering, Austria). In this model, baseline non-feedback data is provided to the model to calibrate the classifier for each subject before the actual classification runs. In addition, the lack of removal of bad trials in our ML approach may explain a difference in the acquired classification accuracies. Also, in Leeuwis et al. (2021) subjects were trained upon online classification, optimizing performance for that specific processing pipeline. Therefore, to make a fair comparison with our DL model, we employed a ML approach using offline classification with no prior training and calibration of the system.

With recent release of large scale EEG datasets (e.g. Cho et al., 2017; Lee et al. 2019), there have been more attempts on employing DL models on signals from large number of participants (e.g., Stieger et al., 2020; Zhang et al., 2021; Ko et al., 2020; Mane et al., 2020), showing the relevance and timeliness of this study in the BCI field. Although these studies report the same conclusion for superiority of the DL approach in MI-BCI classification, their methodology and approach in building the DL model is different from our study. For instance, Stieger et al. (2020) trained a CNN model with high density EEG (64 channel) to classify a 4-class MI task. Mane et al. (2020) and Ko et al. (2020) focused on feature representations in the model; Mane et al. (2020) employed Filter-Bank CNN to decompose data into multiple frequency bands and extract spatially discriminative patterns in each band, and Ko et al. (2020) applied a Multi-Scale Neural Network to exploit spatio-spectral-temporal features for all BCI paradigms. Zhang et al. (2021) focused on transfer learning and employed a CNN model to develop a subject-independent classifier. Therefore, while our study pursues a similar goal, it dissociates itself from past research by conducting a statistically supported subject-wise comparison between the DL and ML approaches and also providing evidence for suitability of the DL approach for inefficient BCI users.

As mentioned before, difference between our study and for example Sakhavi et al. (2015), Gao et al. (2018) and Li et al. (2020) is the employment of pre-processing and feature extraction techniques before applying a DL approach. The only previous study that concentrated on building a CNN model for classification of binary class MI task without implementing any feature engineering technique was conducted by Tang et al. (2017), who achieved 86% mean accuracy for their CNN model. However, they recorded EEG signals from 28 electrodes (as compared to 16 electrodes in this study) and their subject size was only two, which does not provide a suitable representation of the general MI-BCI users. Additionally, Tang et al. (2017) applied 8–30 Hz bandpass filter on the raw data before passing them to the model and the subjects in their study did not receive any feedback during the MI task, which is an important factor in MI learning and online operation of the BCI systems (Alimardani et al., 2016). This makes it difficult to interpret the outcome of Tang et al. (2017) and can perhaps explain the higher accuracy that was achieved by them. In this study, we recruited 54 participants including MI-BCI inefficients (Leeuwis & Alimardani, 2020) and ensured that the participants were learning during the experiment through practice trials and feedback provided by the BCI system.

An important finding of this study was that the CNN model outperformed the CSP+LDA model for all of the subjects, achieving 11-30% accuracy improvement for 92.59% of the subjects. Hence, deducing from the better performance of the DL model compared to ML approach and also from previous studies (Sakhavi et al., 2015; Gao et al., 2018; Li et al., 2020; Tang et al., 2017; Stieger et al., 2020; Zhang et al., 2021; Ko et al., 2020; Mane et al., 2020), it can be concluded that regardless of the users’ ability to generate MI-specific sensorimotor oscillations, CNN models are more effective in extracting intrinsic features from EEG signals and thereon, can perform MI classification with higher accuracy level. This study also revealed that the CNN model was able to capture better understanding of the MI brain patterns in inefficient users than the CSP+LDA model. CNN significantly improved the classification accuracy for those users whose performance was lower when the conventional CSP+LDA model was adopted.

BCI inefficiency has long been seen as a human factor problem in the literature. Only recently, Stieger et al. (2020) suggested that DL approaches might increase accuracies for low aptitude performers, thereby enabling some of them to reach performance above the threshold of 70% accuracy. Our study supports their finding by showing that indeed; the DL approach could significantly improve the classification performance of low performers, supporting the arguments by Thompson (2019), who states that poor performance of training should not be always blamed on the user. Hereby, this study shows that designing an effective classifier using a DL approach could be more reliable in developing robust MI-BCI applications and this also overcomes the issues with BCI inefficiency.

Yet another advantage of the DL approach is that it allows automatic discovery of discriminative features in raw data. Therefore, it is reasonable to consider recording and inclusion of more EEG signals from other brain regions for the model training. Stieger et al. (2020) showed that motor imagery processes might be extended beyond the sensorimotor cortex and mu suppression patterns, indicating that the application of deep learning might be beneficial in extracting such brain activity patterns for inefficient users. Future research can extend our findings by employing a full-scalp recording and showing how this can impact the performance of the CNN model across subjects and thereby future design of more individual-tailored classifiers for all users, especially inefficient users.

The BCI performance is a product of the interplay between the BCI system and the user (Alimardani et al., 2014); therefore, the importance of user training and the ‘*human in the loop*’ cannot be overlooked. Motivation and feedback play an important role in user’s learning of the MI task (Roc et al., 2020, Alimardani et al., 2018). Hence, interaction with a MI-BCI should be established on an engaging platform where the users feel engaged and enjoy the process during experimentation (Roc et al., 2020; Femke et al., 2010). Additionally, detailed instructions on how to perform the mental task of MI should be provided to the users to give them a clear cognitive strategy during BCI training (Roc et al., 2020). This helps to offset the cognitive load on participants and results in stable brain signals, which in turn contributes towards developing an efficient BCI system (Roc et al., 2020). This study employed a classic screen-based feedback bar to provide feedback to the user during data collection. Past studies have shown that embodied feedback in form of virtual or robotic hands can improve interaction between the user and the BCI system (Skola & Liarokapis, 2018; Alimardani et al., 2016). Future studies should attempt to replicate the results of this study with a more engaging and realistic feedback that could lead to generation of more distinguished brain patterns by the user at the data collection stage.

Although this study presents evidence that a DL approach outperforms a ML model for subject-specific classification of the MI task, the question remains whether the proposed CNN model will be able to perform equally well on new subjects who might have different EEG signals. A general challenge in the development and application of MI-BCI systems is their long calibration time (Singh et al., 2019). In order to reduce the calibration time or completely eliminate it, past research has proposed transfer learning in which common information across subjects or sessions is mined and used for training of the classifier to improve the prediction for a new target subject (Azab et al., 2019). However, most transfer learning methodologies focus on extracting features and adapting them from the source subject(s) to the target subject, whereas in DL models with an end-to-end decoding, the neural network itself should be able to do this with little data pre-processing (Zhang et al., 2021). Thus, it becomes important to expand this research in the future with transfer learning methods and evaluate the performance of the proposed CNN model on new targets.

In this study, classification was performed offline. This is not suitable for continuous BCI control where the classifier is constantly updated (Wolpaw & McFarland, 2004), because fluidly controlling an external device is not equal to outputting one command at the end of a trial (Edelman et al., 2019). Stieger et al. (2020) simulated continuous control by providing feedback based on the estimated class output of their CNN every 40 milliseconds and showed that CNN applied on all 64 electrodes made decisions earlier with the threshold degree of confidence and could therefore be applied to make faster decisions in continuous control compared to CNN trained on only motor area electrodes. Their proposal suggests that CNN is applicable for continuous control. Therefore, the accuracy of our classifier providing online continuous feedback should be examined in future research.

In sum, this study aimed to show the potential of DL for MI-EEG classification as opposed to the state-of-the-art ML classifiers. Our results show that compared to the conventional CSP+LDA model, the CNN model, which was trained and tested on raw EEG signals, could achieve significantly higher classification performance for all users, but especially for inefficient users. Applying DL to BCI applications is a burgeoning field, which requires large dataset for development and validation. This study dissociates itself from previous reports by employing a large dataset of 54 subjects and thus sufficiently reflecting the inter-subject variability among BCI users. One of the main advantages of using DL classifier is to eliminate the pre-processing and feature extraction stages used to build an ML classifier. Raw data collected from EEG can directly be fed into a DL classifier. Future studies should be conducted by deploying the proposed CNN model on new subjects to evaluate the performance of the model and to examine whether the same model can be employed for subject-independent classifiers.

## 5 Conclusion

In this research, we evaluated the benefits of DL in improving the performance of motor imagery BCIs. We extracted the performance of a CNN model trained on raw EEG signals from 54 subjects and statistically compared it to that of CSP+LDA, which is a popular ML classifier for binary classification of the MI task. The results revealed that the CNN model significantly outperformed the traditional CSP+LDA classifier by increasing classification accuracy for all 54 subjects in this study.

Moreover, it was shown that the CNN model benefited the inefficient BCI users significantly more than high performers. Thus, we conclude that DL classifiers show promise for future MI-BCI applications for all users as opposed to current state-of-art ML-based BCI systems, which demand extensive effort in pre-processing and feature extraction and yet are impractical for some users. Future studies should further investigate the robustness of the proposed CNN model in real-time MI-BCI applications

## 6 Acknowledgements

Authors would like to thank Alissa Paas for her assistance in collecting the data.

## References

Abbas W, Khan NA. DeepMI: deep learning for multiclass motor imagery classification. In2018 40th Annual International Conference of the IEEE Engineering in Medicine and Biology Society (EMBC) 2018 Jul 18 (pp. 219-222). IEEE. doi: 10.1109/EMBC.2018.8512271

Aggarwal S, Chugh N. Signal processing techniques for motor imagery brain computer interface: A review. Array. 2019 Jan 1;1:100003. doi: 10.1016/j.array.2019.100003

Ahn M, Jun SC. Performance variation in motor imagery brain–computer interface: a brief review. Journal of Neuroscience Methods. 2015 Mar 30;243:103–10. doi: 10.1016/j.jneumeth.2015.01.033

Al-Ani T, Trad D, Somerset VS. Signal processing and classification approaches for brain-computer interface. Intelligent and Biosensors. 2010 Jan 1:25–66. doi: 10.5772/7032

Albawi S, Bayat O, Al-Azawi S, Ucan ON. Social touch gesture recognition using convolutional neural network. Computational Intelligence and Neuroscience. 2018 Oct 8;2018. doi: 10.1155/2018/6973103

Alimardani M, Nishio S, Ishiguro H. Effect of biased feedback on motor imagery learning in BCI-teleoperation system. Frontiers in Systems Neuroscience. 2014 Apr 9;8:52. doi: 10.3389/fnsys.2014.00052

Alimardani M, Nishio S, Ishiguro H. The importance of visual feedback design in BCIs; from embodiment to motor imagery learning. PloS one. 2016 Sep 6;11(9):e0161945. doi: 10.1371/journal.pone.0161945

Alimardani M, Nishio S, Ishiguro H. Brain-computer interface and motor imagery training: The role of visual feedback and embodiment. Evolving BCI Therapy-Engaging Brain State Dynamics. 2018 Oct 17;2:64. doi: 10.5772/intechopen.78695

Allison BZ, Neuper C. Could anyone use a BCI?. In Brain-computer interfaces 2010 (pp. 35–54). Springer, London. doi: 10.1007/978-1-84996-272-8_3

Alom MZ, Taha TM, Yakopcic C, Westberg S, Sidike P, Nasrin MS, Van Esesn BC, Awwal AA, Asari VK. The history began from alexnet: A comprehensive survey on deep learning approaches. arXiv preprint 1803.01164. 2018 Mar 3.

AlZoubi O, Koprinska I, Calvo RA. Classification of brain-computer interface data. In Proceedings of the 7th Australasian Data Mining Conference-Volume 87 2008 Nov 27 (pp. 123–131).

Ang KK, Chin ZY, Wang C, Guan C, Zhang H. Filter bank common spatial pattern algorithm on BCI competition IV datasets 2a and 2b. Frontiers in Neuroscience. 2012 Mar 29;6:39. doi: 10.3389/fnins.2012.00039

Avanzini P, Fabbri-Destro M, Dalla Volta R, Daprati E, Rizzolatti G, Cantalupo G. The dynamics of sensorimotor cortical oscillations during the observation of hand movements: an EEG study. PLoS One. 2012 May 18;7(5):e37534. doi: 10.1371/journal.pone.0037534

Aydemir O, Kayikcioglu T. Comparing common machine learning classifiers in low-dimensional feature vectors for brain computer interface applications. International Journal of Innovative Computing, Information and Control. 2013 Mar;9(3):1145–57.

Azab AM, Mihaylova L, Ang KK, Arvaneh M. Weighted transfer learning for improving motor imagery-based brain–computer interface. IEEE Transactions on Neural Systems and Rehabilitation Engineering. 2019 Jun 17;27(7):1352–9. doi: 10.1109/TNSRE.2019.2923315

Barros ES, Neto N. Classification Procedure for Motor Imagery EEG Data. InInternational Conference on Augmented Cognition 2018 Jul 15 (pp. 201–211). Springer, Cham. doi: 10.1007/978-3-319-91470-1_17

Bashashati H, Ward RK, Birch GE, Bashashati A. Comparing different classifiers in sensory motor brain computer interfaces. PloS one. 2015 Jun 19;10(6):e0129435. doi: 10.1371/journal.pone.0129435

Bharne PP, Kapgate DA. Review of Classification Techniques in Brain Computer Interface. International Journal of Computer Sciences and Engineering. 2014;2:68–72.

Browniee, J. 5 Step Life-Cycle for Neural Network Models in Keras. Machine Learning Mastery. 2016. Retrieved from https://machinelearningmastery.com: https://machinelearningmastery.com/5-step-life-cycle-neural-network-modelskeras/. [Accessed May 1, 2020]

Cho H, Ahn M, Kwon M, Jun SC. A step-by-step tutorial for a motor imagery–based BCI. In Brain–Computer Interfaces Handbook 2018 Jan 9 (pp. 445-460). CRC Press.

Cho H, Ahn M, Ahn S, Kwon M, Jun SC. EEG datasets for motor imagery brain– computer interface. GigaScience. 2017 Jul;6(7):gix034. doi: 10.1093/gigascience/gix034

Craik A, He Y, Contreras-Vidal JL. Deep learning for electroencephalogram (EEG) classification tasks: a review. Journal of Neural Engineering. 2019 Apr 9;16(3):031001. doi: 10.1088/1741-2552/ab0ab5

Dai M, Zheng D, Na R, Wang S, Zhang S. EEG classification of motor imagery using a novel deep learning framework. Sensors. 2019 Jan;19(3):551. doi: 10.3390/s19030551

Dharamsi T, Das P, Pedapati T, Bramble G, Muthusamy V, Samulowitz H, Varshney KR, Rajamanickam Y, Thomas J, Dauwels J. Neurology-as-a-Service for the Developing World. arXiv preprint 1711.06195. 2017 Nov 16.

Edelman BJ, Meng J, Suma D, Zurn C, Nagarajan E, Baxter BS, Cline CC, He B. Noninvasive neuroimaging enhances continuous neural tracking for robotic device control. Science robotics. 2019 Jun 19;4(31). doi: 10.1126/scirobotics.aaw6844

Nijboer F, Birbaumer N, Kubler A. The influence of psychological state and motivation on brain–computer interface performance in patients with amyotrophic lateral sclerosis–a longitudinal study. Frontiers in Neuroscience. 2010 Jul 21;4:55. doi: 10.3389/fnins.2010.00055

Gao G, Shang L, Xiong K, Fang J, Zhang C, Gu X. EEG classification based on sparse representation and deep learning. NeuroQuantology. 2018;16(6). doi: 10.14704/nq.2018.16.6.1666

Goodfellow, I., Bengio, Y., & Courville, A. Deep Learning (Vol. URL (http://www.deeplearningbook.org)). MIT Press. 2016. Retrieved from http://www.deeplearningbook.org. [Accessed April 20, 2019]

Guan S, Zhao K, Yang S. Motor imagery EEG classification based on decision tree framework and Riemannian geometry. Computational Intelligence and Neuroscience. 2019 Jan 21;2019. doi: 10.1155/2019/5627156

Hasan MR, Ibrahimy MI, Motakabber SM, Shahid S. Classification of multichannel EEG signal by linear discriminant analysis. In Progress in Systems Engineering 2015 (pp. 279-282). Springer, Cham.

Hsu WY. EEG-based motor imagery classification using neuro-fuzzy prediction and wavelet fractal features. Journal of Neuroscience Methods. 2010 Jun 15;189(2):295–302. doi: 10.1016/j.jneumeth.2010.03.030

Keras. Convolutional Layers. Keras Documentation. 2019. Retrieved from https://keras.io/layers/convolutional/. [Accessed March 20, 2019].

Khan J, Bhatti MH, Khan UG, Iqbal R. Multiclass EEG motor-imagery classification with sub-band common spatial patterns. EURASIP Journal on Wireless Communications and Networking. 2019 Dec;2019(1):1–9. doi: 10.1186/s13638-019-1497-y

Kingma DP, Ba J. Adam: A method for stochastic optimization. arXiv preprint 1412.6980. 2014 Dec 22.

Ko W, Jeon E, Jeong S, Suk HI. Multi-Scale Neural Network for EEG Representation Learning in BCI. IEEE Computational Intelligence Magazine. 2021 Apr 13;16(2):31–45.

Kosmyna N, Lécuyer A. A conceptual space for EEG-based brain-computer interfaces. PloS one. 2019 Jan 3;14(1):e0210145. doi: 10.1371/journal.pone.0210145

Kumar S, Sharma A, Tsunoda T. An improved discriminative filter bank selection approach for motor imagery EEG signal classification using mutual information. BMC bioinformatics. 2017 Dec;18(16):125–37. doi: 10.1186/s12859-017-1964-6

Kwon OY, Lee MH, Guan C, Lee SW. Subject-independent brain–computer interfaces based on deep convolutional neural networks. IEEE Transactions on Neural Networks and Learning Systems. 2019 Nov 13;31(10):3839–52. doi: 10.1109/TNNLS.2019.2946869

Lebedev MA, Nicolelis MA. Brain-machine interfaces: From basic science to neuroprostheses and neurorehabilitation. Physiological Reviews. 2017 Apr;97(2):767–837. doi: 10.1152/physrev.00027.2016

Lee MH, Kwon OY, Kim YJ, Kim HK, Lee YE, Williamson J, Fazli S, Lee SW. EEG dataset and OpenBMI toolbox for three BCI paradigms: an investigation into BCI illiteracy. GigaScience. 2019 May;8(5):giz002. doi: 10.1093/gigascience/giz002

Leeuwis N, Alimardani M. High Aptitude Motor-Imagery BCI Users Have Better Visuospatial Memory. In2020 IEEE International Conference on Systems, Man, and Cybernetics (SMC) 2020 Oct 11 (pp. 1518-1523). IEEE.

Leeuwis N, Paas A, Alimardani M. Vividness of Visual Imagery and Personality Impact Motor-Imagery Brain Computer Interfaces. Frontiers in Human Neuroscience. 2021;15. doi: 10.3389/fnhum.2021.634748

Li F, He F, Wang F, Zhang D, Xia Y, Li X. A novel simplified convolutional neural network classification algorithm of motor imagery EEG signals based on deep learning. Applied Sciences. 2020 Jan;10(5):1605. doi: 10.3390/app10051605

Lotte F, Bougrain L, Cichocki A, Clerc M, Congedo M, Rakotomamonjy A, Yger F. A review of classification algorithms for EEG-based brain–computer interfaces: a 10 year update. Journal of Neural Engineering. 2018 Apr 16;15(3):031005. doi: 10.1088/1741-2552/aab2f2

Lotte F, Guan C. Regularizing common spatial patterns to improve BCI designs: unified theory and new algorithms. IEEE Transactions on biomedical Engineering. 2010 Sep 30;58(2):355–62. doi: 10.1109/TBME.2010.2082539.

Lotte F, Faller J, Guger C, Renard Y, Pfurtscheller G, Lécuyer A, Leeb R. Combining BCI with virtual reality: towards new applications and improved BCI. InTowards Practical Brain-Computer Interfaces 2012 (pp. 197–220). Springer, Berlin, Heidelberg.

Mane R, Robinson N, Vinod AP, Lee SW, Guan C. A Multi-view CNN with Novel Variance Layer for Motor Imagery Brain Computer Interface. In2020 42nd Annual International Conference of the IEEE Engineering in Medicine & Biology Society (EMBC) 2020 Jul 20 (pp. 2950-2953). IEEE. doi: 10.1109/EMBC44109.2020.9175874

Marchesotti S, Bassolino M, Serino A, Bleuler H, Blanke O. Quantifying the role of motor imagery in brain-machine interfaces. Scientific Reports. 2016 Apr 7;6(1):1–2. doi: 10.1038/srep24076

McFarland DJ, Wolpaw JR. Sensorimotor rhythm-based brain-computer interface (BCI): feature selection by regression improves performance. IEEE Transactions on Neural Systems and Rehabilitation Engineering. 2005 Sep 12;13(3):372–9. doi: 10.1109/TNSRE.2005.848627

Miao M, Hu W, Yin H, Zhang K. Spatial-frequency feature learning and classification of motor imagery EEG based on deep convolution neural network. Computational and Mathematical Methods in Medicine. 2020 Jul 20;2020. doi: 10.1155/2020/1981728

Millán JD, Rupp R, Mueller-Putz G, Murray-Smith R, Giugliemma C, Tangermann M, Vidaurre C, Cincotti F, Kubler A, Leeb R, Neuper C. Combining brain– computer interfaces and assistive technologies: state-of-the-art and challenges. Frontiers in Neuroscience. 2010 Sep 7;4:161. doi: 10.3389/fnins.2010.00161

Müller KR, Krauledat M, Dornhege G, Curio G, Blankertz B. Machine learning techniques for brain-computer interfaces. Biomed. Tech. 2004 Dec;49(1):11–22.

Nicolas-Alonso LF, Gomez-Gil J. Brain Computer Interfaces, a Review. Sensors. 2012 Feb;12(2):1211–79. doi: 10.3390/s120201211

Park Y, Chung W. Selective feature generation method based on time domain parameters and correlation coefficients for Filter-Bank-CSP BCI systems. Sensors. 2019 Jan;19(17):3769. doi: 10.3390/s19173769

Pfurtscheller G, Brunner C, Schlögl A, Da Silva FL. Mu rhythm (de) synchronization and EEG single-trial classification of different motor imagery tasks. NeuroImage. 2006 May 15;31(1):153–9.

Pfurtscheller G, Neuper C. Motor imagery and direct brain-computer communication. Proceedings of the IEEE. 2001 Jul;89(7):1123–34. doi: 10.1109/5.939829

Qiao W, Bi X. Deep spatial-temporal neural network for classification of EEG-based motor imagery. InProceedings of the 2019 International Conference on Artificial Intelligence and Computer Science 2019 Jul 12 (pp. 265–272). doi: 10.1145/3349341.3349414

Radiuk PM. Impact of training set batch size on the performance of convolutional neural networks for diverse datasets. Information Technology and Management Science. 2017 Dec 20;20(1):20–4. doi: 10.1515/itms-2017-0003

Raza H, Cecotti H, Prasad G. Optimising frequency band selection with forward-addition and backward-elimination algorithms in EEG-based brain-computer interfaces. In2015 international joint conference on neural networks (IJCNN) 2015 Jul 12 (pp. 1-7). IEEE. doi: 10.1109/IJCNN.2015.7280737

Robinson N, Lee SW, Guan C. EEG representation in deep convolutional neural networks for classification of motor imagery. In2019 IEEE International Conference on Systems, Man and Cybernetics (SMC) 2019 Oct 6 (pp. 1322-1326). IEEE.

Roc A, Pillette L, Mladenovic J, Benaroch C, N’Kaoua B, Jeunet C, Lotte F. A review of user training methods in brain computer interfaces based on mental tasks. Journal of Neural Engineering. 2020 Nov 12. doi: 10.1088/1741-2552/abca17

Roy Y, Banville H, Albuquerque I, Gramfort A, Falk TH, Faubert J. Deep learning-based electroencephalography analysis: a systematic review. Journal of Neural Engineering. 2019 Aug 14;16(5):051001. doi: 10.1088/1741-2552/ab260c

Saha S, Baumert M. Intra-and inter-subject variability in EEG-based sensorimotor brain computer interface: a review. Frontiers in computational neuroscience. 2020 Jan 21;13:87. doi: 10.3389/fncom.2019.00087

Sakhavi S, Guan C, Yan S. Parallel convolutional-linear neural network for motor imagery classification. In2015 23rd European Signal Processing Conference (EUSIPCO) 2015 Aug 31 (pp. 2736-2740). IEEE. doi: 10.1109/EUSIPCO.2015.7362882

Schirrmeister RT, Springenberg JT, Fiederer LD, Glasstetter M, Eggensperger K, Tangermann M, Hutter F, Burgard W, Ball T. Deep learning with convolutional neural networks for EEG decoding and visualization. Human brain mapping. 2017 Nov;38(11):5391–420. doi: 10.1002/hbm.23730

Selim S, Tantawi MM, Shedeed HA, Badr A. A CSP\AM-BA-SVM Approach for Motor Imagery BCI System. IEEE Access. 2018 Aug 31;6:49192–208. doi: 10.1109/ACCESS.2018.2868178

Sharma, S. Epoch vs Batch Size vs Iterations. 2017, September 27. Retrieved from towards data science: https://towardsdatascience.com/epoch-vs-iterations-vs-batch-size4dfb9c7ce9c9. [Accessed May 15, 2020]

Shashibala T, Gawali BW. Brain computer interface applications and classification techniques. International Journal of Engineering and Computer Science. 2016;5(7):17260–7.

Shen Y, Lu H, Jia J. Classification of motor imagery EEG signals with deep learning models. InInternational Conference on Intelligent Science and Big Data Engineering 2017 Sep 22 (pp. 181–190). Springer, Cham. doi: 10.1007/978-3-319-67777-4_16

Singh A, Lal S, Guesgen HW. Reduce calibration time in motor imagery using spatially regularized symmetric positives-definite matrices based classification. Sensors. 2019 Jan;19(2):379. doi: 10.3390/s19020379

Škola F, Liarokapis F. Embodied VR environment facilitates motor imagery brain– computer interface training. Computers & Graphics. 2018 Oct 1;75:59–71. doi: 10.1016/j.cag.2018.05.024

Stieger J, Engel S, Suma D, He B. Benefits of deep learning classification of continuous noninvasive brain-computer interface control. Journal of Neural Engineering. 2021 May 26. doi: 0.1016/j.cag.2018.05.024

Tang Z, Li C, Sun S. Single-trial EEG classification of motor imagery using deep convolutional neural networks. Optik. 2017 Feb 1;130:11–8. doi: 10.1016/j.ijleo.2016.10.117

Tayeb Z, Fedjaev J, Ghaboosi N, Richter C, Everding L, Qu X, Wu Y, Cheng G, Conradt J. Validating deep neural networks for online decoding of motor imagery movements from EEG signals. Sensors. 2019 Jan;19(1):210. doi: 10.3390/s19010210

TensorFlow. Overfit and underfit. 2020, July 10. Retrieved from TensorFlow: https://www.tensorflow.org/tutorials/keras/overfit_and_underfit. [Accessed April 02, 2020]

Thompson MC. Critiquing the concept of BCI illiteracy. Science and engineering ethics. 2019 Aug;25(4):1217–33. doi: 10.1007/s11948-018-0061-1

Torres, J. Learning Process of a Neural Network. How Do Artificial Neural Networks Learn? 2020, April 21. Retrieved from towards data science: https://towardsdatascience.com/learning-process-of-a-deep-neural-network-5a9768d7a651. [Accessed May 05, 2020].

Ullah I, Manzo M, Shah M, Madden M. Graph Convolutional Networks: analysis, improvements and results. arXiv preprint 1912.09592. 2019 Dec 19.

Vidaurre C, Kawanabe M, von Bünau P, Blankertz B, Müller KR. Toward unsupervised adaptation of LDA for brain–computer interfaces. IEEE Transactions on Biomedical Engineering. 2010 Nov 18;58(3):587–97. doi: 10.1371/journal.pone.0123727

Wang P, Jiang A, Liu X, Shang J, Zhang L. LSTM-based EEG classification in motor imagery tasks. IEEE transactions on neural systems and rehabilitation engineering. 2018 Oct 18;26(11):2086–95. doi: 10.1109/TNSRE.2018.2876129

Wang Z, Ma Z, Du X, Dong Y, Liu W. Research on the Key Technologies of Motor Imagery EEG Signal Based on Deep Learning. Journal of Autonomous Intelligence. 2019;2(4):1–4. doi: 10.32629/jai.v2i4.60

Wierzgała P, Zapała D, Wojcik GM, Masiak J. Most popular signal processing methods in motor-imagery BCI: a review and meta-analysis. Frontiers in Neuroinformatics. 2018 Nov 6;12:78. doi: 10.3389/fninf.2018.00078

Wolpaw JR, McFarland DJ. Control of a two-dimensional movement signal by a noninvasive brain-computer interface in humans. Proceedings of the National Academy of Sciences. 2004 Dec 21;101(51):17849–54. doi: 10.1073/pnas.0403504101

Yahya N, Musa H, Ong ZY, Elamvazuthi I. Classification of motor functions from electroencephalogram (EEG) signals based on an integrated method comprised of common spatial pattern and wavelet transform framework. Sensors. 2019 Jan;19(22):4878. doi: 10.3390/s19224878

Yang H, Sakhavi S, Ang KK, Guan C. On the use of convolutional neural networks and augmented CSP features for multi-class motor imagery of EEG signals classification. In2015 37th Annual International Conference of the IEEE Engineering in Medicine and Biology Society (EMBC) 2015 Aug 25 (pp. 2620-2623). IEEE. doi: 10.1109/EMBC.2015.7318929

Yuksel A, Olmez T. A neural network-based optimal spatial filter design method for motor imagery classification. PloS one. 2015 May 1;10(5):e0125039. doi: 10.1371/journal.pone.0125039

Zapała D, Zabielska-Mendyk E, Augustynowicz P, Cudo A, Jaśkiewicz M, Szewczyk M, Kopiś N, Francuz P. The effects of handedness on sensorimotor rhythm desynchronization and motor-imagery BCI control. Scientific Reports. 2020 Feb 7;10(1):1–1. doi: 10.1038/s41598-020-59222-w

Pérez Zapata AF. Classification of Motor Imagery EEG Signals Using a CNN Architecture and a Meta-heuristic Optimization Algorithm for Selecting Training Parameters.

Zhang K, Robinson N, Lee SW, Guan C. Adaptive transfer learning for EEG motor imagery classification with deep Convolutional Neural Network. Neural Networks. 2021 Apr 1;136:1–0. doi: 10.1016/j.neunet.2020.12.013

